# Short term changes in the proteome of human cerebral organoids induced by 5-methoxy-N,N-dimethyltryptamine

**DOI:** 10.1101/108159

**Authors:** Vanja Dakic, Juliana Minardi Nascimento, Rafaela Costa Sartore, Renata de Moraes Maciel, Draulio B. de Araujo, Sidarta Ribeiro, Daniel Martins-de-Souza, Stevens Rehen

**Affiliations:** D’Or Institute for Research and Education (IDOR), Rio de Janeiro, Brazil; Institute of Biomedical Sciences, Federal University of Rio de Janeiro, Rio de Janeiro, Brazil; Laboratory of Neuroproteomics, Institute of Biology, Department of Biochemistry and Tissue Biology, University of Campinas (UNICAMP), Campinas, Brazil; Brain Institute, Federal University of Rio Grande do Norte, Natal, Brazil

**Author notes:** These authors contributed equally to this work (Vanja Dakic, Juliana Minardi Nascimento).

**Keywords:** NPC, organoid, 5-MeO-DMT, plasticity, proteomics

## Abstract

Dimethyltryptamines are hallucinogenic serotonin-like molecules present in traditional Amerindian medicine (e.g. *Ayahuasca)* recently associated with cognitive gains, antidepressant effects and changes in brain areas related to attention. Historical and technical restrictions impaired understanding how such substances impact human brain metabolism. Here we used shotgun mass spectrometry to explore proteomic differences induced by 5-methoxy-N,N-dimethyltryptamine (5-MeO-DMT) on human cerebral organoids. Out of the 6,728 identified proteins, 934 were found differentially expressed in 5-MeO-DMT-treated cerebral organoids. *In silico* systems biology analyses support 5-MeO-DMT’s anti-inflammatory effects and reveal a modulation of proteins associated with long-term potentiation, the formation of dendritic spines, including proteins involved in cellular protrusion formation, microtubule dynamics and cytoskeletal reorganization. These results offer possible mechanistic insights into the neuropsychological changes caused by the ingestion of substances rich in dimethyltryptamines.

## Introduction

Dimethyltryptamines are naturally occurring hallucinogenic molecules hypothesized to be involved in spontaneous altered states of consciousness, such as dreams, free imagination and insightful creativity (Barker *et al,* 2012; Strassman, 2001). N,N-dimethyltryptamine (N,N-DMT) and bufotenine (5-HO-DMT) have been traditionally used as entheogens by Amerindians (McKenna, 2004; Ott, 2001) as major active ingredients of a brew called *Ayahuasca* and the *Virola* snuff (Holmstedt and Lindgren, 1967). The popularity of *Ayahuasca* as part of religious ceremonies continues to spread in South America and other countries (Labate and Feeney, 2012), possibly motivated by its strong antidepressant effects (Osório Fde *et al,* 2015; Sanches *et al,* 2016). Chronic *Ayahuasca* ingestion has been associated with cognitive gains and structural brain changes in areas related to attention, self-referential thought, and internal mentation (Bouso *et al,* 2012, 2015).

The search for the molecular mechanisms underlying the effects of dimethyltryptamines showed that N,N-DMT and 5-methoxy-N,N-dimethyltryptamine (5-MeO-DMT), two closely related metabolic products, can act as systemic endogenous regulators of inflammation and immune homeostasis through both 5-hydroxytryptamine receptors (5-HTRs) and sigma 1 receptors (σ-1Rs) (Fontanilla *et al,* 2009; Szabo *et al,* 2014). Under severe hypoxia, N,N-DMT robustly increased the survival of cultured human cortical neurons *in vitro,* monocyte-derived macrophages, and dendritic cells acting through σ-1Rs (Szabo *et al,* 2016). The direct evidence of neuro-immune communication and neuroregenerative effects of N,N-DMT and 5-MeO-DMT greatly enhanced expectations for dimethyltryptamine research.

Our limited understanding of the physiological activity of dimethyltryptamine and other classic psychedelic substances is caused by legal restrictions on such research (Nutt *et al,* 2013) but also by the lack of adequate experimental models (Hanks and González-Maeso, 2013; de la Torre and Farré, 2004; Vollenweider and Kometer, 2010). In the past few years, considerable progress has been made regarding the neural differentiation of human pluripotent stem cells into mature neurons and cerebral organoids (Kelava and Lancaster, 2016). Human neural progenitors (hNPC) are useful cell systems for high-throughput screening due to their homogeneity, with little complexity and limited differentiation potential. On the other hand, cerebral organoids are complex three-dimensional (3D) culture systems with multiple cell types that self-organize into various brain regions similarly to those *in vivo,* including the cerebral cortex, ventral forebrain, midbrain-hindbrain boundary, and hippocampus (Lancaster and Knoblich, 2014; Qian *et al,* 2016). A comparison of gene expression programs of human fetal neocortex and *in vitro* cortical development by single-cell RNA sequencing found remarkable similarities (Camp *et al,* 2015). Cerebral organoids are being widely applied to human-specific biological questions and purposes, since they have been proved to be a good system for drug testing. They may well recapitulate effects in the human nervous system, particularly related to plasticity and growth (Garcez *et al,* 2016; Lancaster and Knoblich, 2014), and they circumvent problems of discrepancies in metabolic pathways occurring in translational studies involving animals. The development of such model offers an exciting new range of opportunities to investigate the molecular responses of human neural tissue to psychoactive substances.

Here we analyzed the effect of 5-MeO-DMT on neural cells and human brain organoids. By employing mass spectrometry-based proteomics to analyze cerebral organoids, we managed to investigate effects in a large scale and in an unbiased manner, and to get insights into the molecular mechanisms and biochemical pathways involved (Martins-de-Souza, 2012). Our results show that 5-MeO-DMT modulates several proteins involved in long-term potentiation (LTP), in addition to morphogenesis and maturation of dendritic spines, while inhibiting neurodegeneration and cell death.

## Materials and methods

### Human embryonic stem cells

BR1 lineage of human embryonic stem cells (hESCs) (Fraga *et al,* 2011) was cultured in mTeSR1 media (Stemcell Technologies) on Matrigel (BD Biosciences) - coated surface. The colonies were manually passaged every seven days and maintained at 37°C in humidified air with 5% CO_2_.

### Human neural progenitor cells

To induce hESCs towards neural differentiation, we adapted Baharvand and coworkers’ protocol (Baharvand *et al*, 2007; Dakic *et al,* 2016). Briefly, 70% confluent human embryonic stem cells were differentiated to the neural lineage in defined adherent culture by retinoic acid and basic fibroblast growth
factor (bFGF) within 18 days of culture. On the 18th day, neural tube-like structures were collected and plated on dishes coated with 10 μg/mL of poly-L-ornithine and 2.5 μg/mL of laminin (Thermo Fisher Scientific). The population of human neural progenitor cells (hNPCs) that migrated from neural tube-like structures was tested for the expression of neuronal markers and expanded. Expansion was done in N2B27 medium supplemented with 25 ng/mL bFGF and 20 ng/mL EGF (Thermo Fisher Scientific). N2B27 medium consisted of DMEM/F-12 supplemented with 1X N2, 1X B-27, 1% penicillin/streptomycin (Thermo Fisher Scientific). Cells were incubated at 37°C and 5% CO_2_. Medium was replaced every other day. hNPCs were expanded for no more than 5 passages. Basic characterization of this culture was published in Dakic et al., 2016.

### High Content Screening

Cell proliferation, cell death and arborization experiments were performed in a High Content Screening (HCS) format. hNPCs (1,500 cells per well) were plated on a 384-multiwell μClear plate (Greiner Bio-One, Kremsmünster, Austria) coated with 100 μg/mL poly-L-ornithine and 10 μg/mL laminin (Thermo Fisher Scientific). After 24h, cells were treated for 4 days in quintuplicate (five wells per condition) with 5-MeO-DMT (Sigma-Aldrich) in N2B27 medium supplemented with bFGF and EGF. Cells were labeled with 10 μM EdU for 2h (cell proliferation) for 30 min prior to fixation and image acquisition. For cell death assessment, cells were labeled with LIVE/DEAD^®^ viability/cytotoxicity kit (Thermo Fisher Scientific). This kit contains two probes: calcein AM and ethidium homodimer (EthD-1). The first one allows measuring of intracellular esterase activity and second one plasma membrane integrity. Mix of probes was done in DMEM/F-12 (without phenol red, Life Technologies), together with the cell-permeant nuclear dye Hoechst. After incubation for 30 min at 37°C and 5% CO_2_, the dye cocktail was replaced by new medium and image acquired. For arborization experiments, neural differentiation was induced 24h after plating by removal of bFGF and EGF from N2B27 medium. Treatment with 5-MeO-DMT was done concomitantly with neural differentiation. Medium was changed after 4 days of treatment and cells were allowed to differentiate for 3 more days. On day 7 cells were fixed for immunocytochemistry.

### High Content Analysis

All images were acquired on Operetta high-content imaging system (Perkin Elmer, USA). For proliferation, incorporated EdU was detected with Alexa Fluor 488 using Click-iT EdU kit (C10351, Invitrogen, Carlsbad, USA) following the manufacturer’s instruction. Total number of cells was calculated by nuclei stained with 1 mg/mL of DAPI (4’,6-diamidino-2-phenylindole). S phase was determined by percentage of total cells labeled with EdU. Images were acquired with a 10x objective with high numerical aperture (NA).

Live cell imaging was performed with LIVE/DEAD^®^ viability/cytotoxicity kit, using temperature and CO_2_ control option (TCO) of Operetta, set to 37°C and 5% CO_2_ at 10x magnification. Quantification analyses were normalized to the number of cells in the well segmented by nucleus dyes.

Neuronal arborization was evaluated on fixed cells stained for MAP2 after 7 days of differentiation. The images were analyzed using the Neurite Outgrowth script of the Harmony software. Briefly, neurites were detected on MAP2 positive cells using the Find Neurite building block, which provides a dedicated algorithm for segmenting neurites. Morphological characteristics of neuronal arborization, such as total neurite length (sum of the length of all neurites attached to the cell), number of extremities, number of segments and number of nodes type I were defined based on selected threshold parameters of the Find Neurite building block.

All analysis sequences were designated by combining segmentation steps with morphological and fluorescence based object characterizations using the image analysis software Harmony 3.5.1 (Perkin Elmer, Waltham, MA, USA).

### Differentiation into cerebral organoids

Differentiation of hESCs into cerebral organoids was based on previously described protocol (Lancaster *et al,* 2013; Sartore *et al,* 2017). Briefly, hESC cells were inoculated into a spinner flask, and to enable embryoid body formation, after six days the medium was changed to neural induction media (DMEM/F12, 1x N2 supplement (Gibco), 2 mM Glutamax (Invitrogen), 1% MEM-NEAA and 1 μg/mL heparin (Sigma) and the aggregates were cultured for five more days. After being embedded in matrigel, differentiation media composed of 1:1 DMEM/F12: Neurobasal (Gibco), 0.5 x N2, 1x B27 minus vitamin A (Gibco), 2 mM Glutamax, 0.5% MEM-NEAA, 0.2 μM 2-Mercaptoethanol and 2.5 μg/mL insulin was used. After 4 days cell aggregates were grown in neuronal differentiation media, composed as aforementioned except by replacing with B27 containing vitamin A (Gibco). The medium was changed once per week. Cerebral organoids were grown for 45 days (30 days in neuronal differentiation media) and the basic characterization of this method was published in Sartore at al., 2017.

### RNA Isolation and PCR Analysis

Total RNA was isolated using the GeneJET RNA Purification Kit (Thermo Scientific) and digested with DNase using DNase I (Invitrogen), following the manufacturer’s instructions. Complementary DNA was generated from 1 μg total RNA using M-MLV Reverse Transcriptase (Invitrogen), according to the manufacturer’s recommendations. PCR was performed using the following primer sequences: GFAP-For: 5’-TTC GAC AGT CAG CCG CAT C-3’ GFAP-Rev: 5’-GAC TCC ACG ACG TAC TCA GC −3’, Sigma receptor 1-For: 5’-AGT AGG ACC ATG CAC TCA CAC C-3’ Sigma receptor 1-Rev: 5’-CCC CAT CCT TAA CTC TAG AAC C −3’, 5-HT_2A_-For: 5’-TTG GGC TAC AGG ACG ATT −3’ 5-HT_2A_-Rev: 5’-GAA GAA AGG GCA CCA CAT C −3’, 5-HT_2C_-For: 5’-TGT CCC TAG CCA TTG CTG ATA TGC −3’ 5-HT2C-Rev: 5’-GCA ATC TTC ATG ATG GCC TTA GTC −3’. Each PCR reaction was carried out for 40 cycles in a reaction mixture containing 0.25 U Taq DNA Polymerase (Invitrogen), 1× Taq DNA Polymerase Buffer containing 1.5 mM MgCl_2_ (Invitrogen), 200 nM of each primer (forward and reverse), 200 μM dNTP mixture containing the four deoxyribonucleotides (dATP, dCTP, dTTP, dGTP), and 0.5 μl of cDNA.

### Immunohistochemistry

On the 45th day of differentiation, cerebral organoids were fixed in 4% paraformaldehyde, incubated with sucrose solutions (10, 20 and 30%) in phosphate buffered saline (PBS), embedded in optimal cutting temperature compound (OCT) and frozen in liquid nitrogen. The organoids were sectioned with a cryostat into 20 μm thick sections. Immunofluorescence was performed using the primary antibodies: anti-MAP2 (M1406, Sigma-Aldrich), anti-AMPAR1 (Abcam, ab86141), anti-NMDAR1 (Abcam, ab28669), anti-sigma receptor 1 (sc-137075, Santa Cruz), anti-5-HT_2A_ (RA24288, Neuromics). Secondary antibodies were used as follows: Alexa Fluor 488 Goat anti-mouse (A11001, Invitrogen) and Alexa Fluor 594 Goat anti-mouse (A-11008, Invitrogen). DAPI was used for nuclei staining. Images were acquired using an Operetta Imaging System (Perkin Elmer) and a Leica TCS SP8 confocal microscope, when specified.

### Treatment of cerebral organoids with 5-MeO-DMT

On day 45 of differentiation, four to five organoids per group were transferred from the spinner flask to a non-adherent dish and treated with 13 μM 5-MeO-DMT (Sigma-Aldrich), 0.3% ethanol (vehicle) or only medium (control), for 24 hours. After treatment, cerebral organoids were pelleted and homogenized in buffer containing 7 M Urea, 2 M thiourea, 4% CHAPS, 70 mM DTT, and Complete Protease Inhibitor Cocktail (Roche) (Maccarrone *et al,* 2014). The homogenates were kept on ice for about 20 min and frozen at −80°C until sample processing for mass spectrometry-based label-free shotgun proteomics. The experiment was repeated three times with the three different derivations of cerebral organoids.

### Sample preparation

Sample lysates were thawed and centrifuged at 10,000 x g for 10 min at 4 °C. The supernatant was collected and subjected to quantification by Qubit^®^ 3.0 Fluorometer (Thermo Fisher Scientific). Each sample (50μg) was subjected to a SDS-PAGE gel electrophoresis. Gel lanes were sliced and digested *in gel* overnight as previously described (Maccarrone *et al,* 2014). Generated peptides were dried in a SpeedVac concentrator and stored at −80°C prior to shotgun mass spectrometry analyses.

### Liquid chromatography-mass spectrometry

Qualitative and quantitative proteomic analysis were performed on a 2D-LC-MS/MS system with ion-mobility-enhanced data-independent acquisitions (Souza *et al,* 2017). Peptides were injected into a two-dimensional reverse phase liquid chromatography using an Acquity UPLC M-Class System (Waters Corporation, Milford, MA) coupled to a Synapt G2-Si mass spectrometer (Waters Corporation, Milford, MA).

In first dimension chromatography, peptides (5 μg) were loaded into a M-Class BEH C18 Column (130 Å, 5 μm, 300 μm X 50 mm, Waters Corporation, Milford, MA). Fractionation was performed using discontinuous steps of acetonitrile (11%, 14%, 17%, 20% and 50% acetonitrile). After each step, peptide loads went for second dimension separation, at nanoACQUITY UPLC HSS T3 Column (100Å, 1.8 μm, 75 μm X 150mm, Waters Corporation, Milford, MA). Peptide elution was achieved by using an acetonitrile gradient from 7% to 40% (v/v) for 54 min at a flow rate of 0.4 μL /min directly into a Synapt G2-Si. The mass spectrometer acquired in a data-independent mode (DIA) with ion mobility separation. This approach called high-definition data-independent mass spectrometry (HDMS^E^) enhances the proteome coverage significantly (Distler *et al,* 2014). MS/MS analyses were performed by nanoelectrospray ionization in positive ion mode nanoESI (+) and a NanoLock Spray (Waters, Manchester, UK) ionization source. The lock mass channel was sampled every 30 s. The mass spectrometer was calibrated with an MS/MS spectrum of [Glu1]-Fibrinopeptide B human (Glu-Fib) solution that was delivered through the reference sprayer of the NanoLock Spray source. Samples were all run in technical and biological triplicates.

### Database search and quantification

Raw data was processed with Progenesis^®^ QI version 2.1 (Waters) and proteins were identified. Quantitative data was processed using dedicated algorithms and searching against the Uniprot human proteomics database (version 2015/09), with the default parameters for ion accounting and quantitation (Li *et al,* 2009). The databases used were reversed “on the fly” during the database queries and appended to the original database to assess the false-positive identification rate. The following parameters were considered in identifying peptides: 1) Digestion by trypsin with at most one missed cleavage; 2) variable modifications by oxidation (M) and fixed modification by carbamidomethyl (C); 3) false discovery rate (FDR) less than 1%. Identifications that did not satisfy these criteria were not considered.

### *In silico* analysis

Protein networks and canonical pathways associated with differentially expressed proteins were identified using software Ingenuity Pathway Analysis (IPA, Ingenuity Systems, Qiagen, Redwood, CA, USA; www.ingenuity.com). This software uses curated connectivity information from the literature to determine interaction network among the differentially expressed proteins and canonical pathways in which they are involved. Here, we have considered information from nervous system tissues and cells, immune cells and stem cells. The significant biological functions are based on Fisher’s exact test. Multiple correlation hypothesis are based on Benjamini-Hochberg (B-H) approach using 1% FDR threshold, the significance of the IPA test is expressed as p-values (Schubert *et al,* 2015).

## Results

### Human neural progenitor cells are unaffected by 5-MeO-DMT

First, we examined the effects of 5-MeO-DMT in hNPCs (detailed characterization in (Dakic *et al,* 2016). hNPCs showed basal expression of σ-1Rs but not 5-HT_2A_ and 5-HT_2C_ (Fig. 1A and B). Using a high content screening analysis, we tested the effects of 5-MeO-DMT (23 nM to 7.11 μΜ) upon cell death, proliferation and differentiation of hNPCs. There was no evidence of change in cell death or proliferation in response to 5-MeO-DMT (Fig. 1C and D). In addition, by quantifying some aspects of dendritic branch complexity, we measured neural arborization based on MAP2 staining of young neurons exposed or not to 5-MeO-DMT. Despite a slight trend, there were no statistically significant differences in the measured parameters (Fig. 1 E, F, G and H).

**Figure 1.**
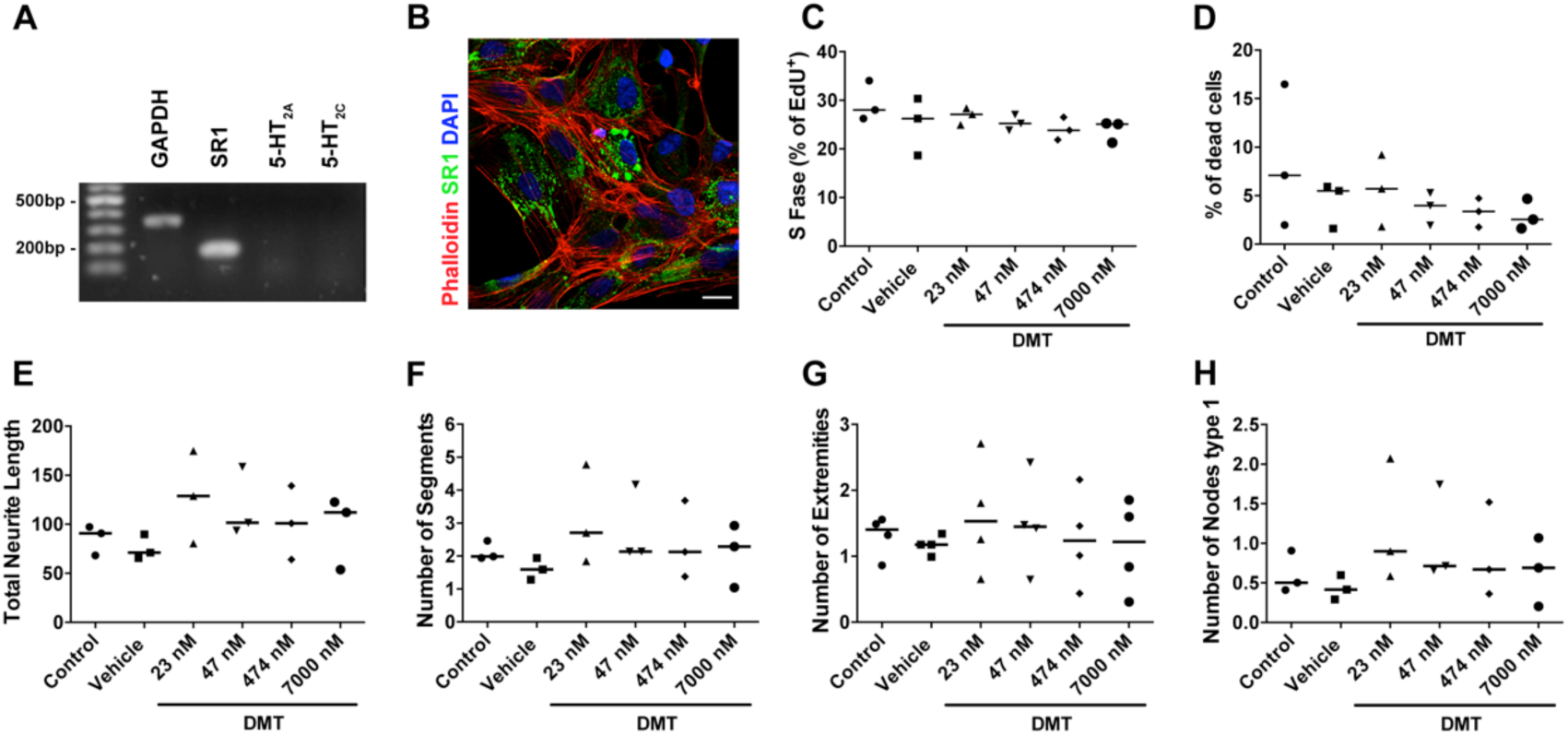
Effects of 5-MeO-DMT on hNPCs. **(A)** Expression of mRNA for internal control (GAPDH), SR1, 5-HT_2A_ and 5-HT_2C_ in hNPCs. **(B)** Confirmation of σ-1R protein (green) expression by immunocytochemistry, phalloidin showing the cytoskeleton (red) and DAPI staining nuclei (blue), scale bar 20 μm. **(C)** Quantification of cell proliferation based on EdU staining after treatment with 5-MeO-DMT. **(D)** Percentage of dead cells in hNPCs treated with 5-MeO-DMT. **(E, F, G, H)** Effects of 5-MeO-DMT on neuronal arborization by quantification of **(E)** total neurite length (sum of the length of all neurites attached to the cell), **(F)** number of segments, **(G)** number of extremities, and **(H)** number of nodes type 1. Bar represents median. Data analyzed by one-way ANOVA with Tukey’s multiple comparison test.

### Human cerebral organoids express 5-MeO-DMT receptors

The lack of alterations in cell death, proliferation or differentiation/arborization could be due to the low cellular diversity and lack of complex interactions among different cell types. Thus, we challenged human cerebral organoids, which recapitulate better the complexity and function of *in vivo* neural circuitry. Basal immunostaining of ionotropic receptors AMPA and NMDA, characteristic of glutamatergic synapses, along with the neuronal marker MAP2 was observed in 45-day-old cerebral organoids (Fig. 2A-D), as previously described (Sartore *et al,* 2017). Glial cells (GFAP+) are also present in organoids, as shown in Fig. 2E. Interestingly, in contrast with hNPCs, we were able to detect the expression of 5-HT2A via PCR and/or immunostaining, as well as of σ-1Rs, the primary pharmacological molecular targets for 5-MeO-DMT. As shown in Figure 2F-G, cells with the 5-HT_2A_ receptor are present within the cerebral organoid, and the sigma-1 receptor was detected as well. RT-PCR confirmed the expression of 5-HT_2A_ and σ-1 receptors, and allowed the detection of serotonin 5-HT_2C_ receptors as well (Fig. 2H). Taken together, these data validate the cerebral organoids as an appropriate platform to seek for 5-MeO-DMT effects in an amenable yet realistic human neuronal network.

**Figure 2.**
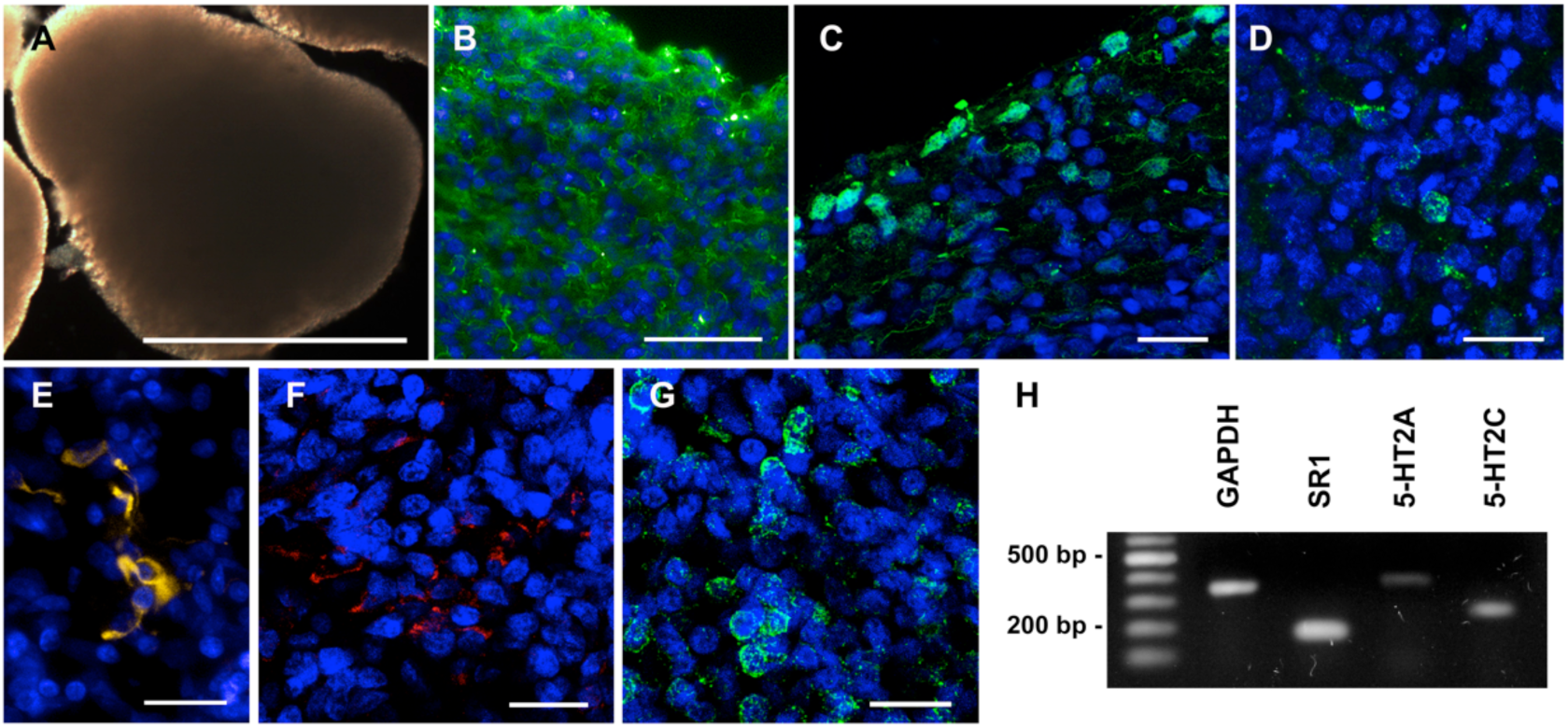
Cerebral organoids express 5-MeO-DMT receptors and different cell type markers. **(A)** Cerebral organoids presenting smooth texture and homogeneous coloring at 45 days of differentiation (scale bar 1000 μm). **(B)** Cerebral organoids are composed by several cell types, including mature neurons, as shown by MAP2 staining. **(C)** Cells expressing AMPAR1 are found in the organoid edge, while **(D)** cells expressing NMDAR1 and **(E)** GFAP are detected within the organoid. **(F)** Cells positive for 5-HT_2A_ receptor, and **(G)** σ-1R, the primary molecular targets for 5-MeO-DMT, are also found in the organoid. Scale bars: A = 1000 μm; B = 50 μm; C, D, E, F, and G = 20 μm. **(H)** The expression of molecular targets for 5-MeO-DMT was also confirmed by RT-PCR.

### 5-MeO-DMT alters the proteome of human cerebral organoids

Due to the complexity of the organoid system, we decided to cast a much wider net to detect potentially important 5-MeO-DMT effects. By analyzing the proteome of organoids with or without treatment, we were able to look for changes in the expression of a considerable number of proteins, in an unbiased approach. Thus, to resolve the proteome of human neural tissue under the effect of 5-MeO-DMT, we analyzed 45-day-old cerebral organoids after 24-hour treatment (Fig. 3A). A total of 144,700 peptides were identified at a false discovery rate (FDR) below 1%. These led to the identification of 6,728 unique proteins by, at least, two unique peptides present in no less than two out of three biological replicates analyzed. Notably, there was an overlap of 99% of identified proteins among all treatment groups (Fig. 3B), demonstrating the robustness of the method. From these commonly identified proteins, we found 934 differentially expressed (using a −2 < Log_2_ ratio > 2 cut-off), comprising 360 downregulated and 574 upregulated proteins, when comparing 5-MeO-DMT and vehicle groups. Functional enrichment for combined up- and downregulated proteins predicted the biological functions of those changes. Regarding diseases or functions, using prediction effect analysis (−2 < z-score > 2.0 is significant for inhibition/activation) (Fig. 3C), we observed a significant activation score for dendritic spine and cellular protrusion formation, microtubule and cytoskeletal organization, and also mild activation of T lymphocyte differentiation. On the other hand, biological functions such as neurodegeneration, cell death, and brain lesion were predicted to be inhibited.

**Figure 3.**
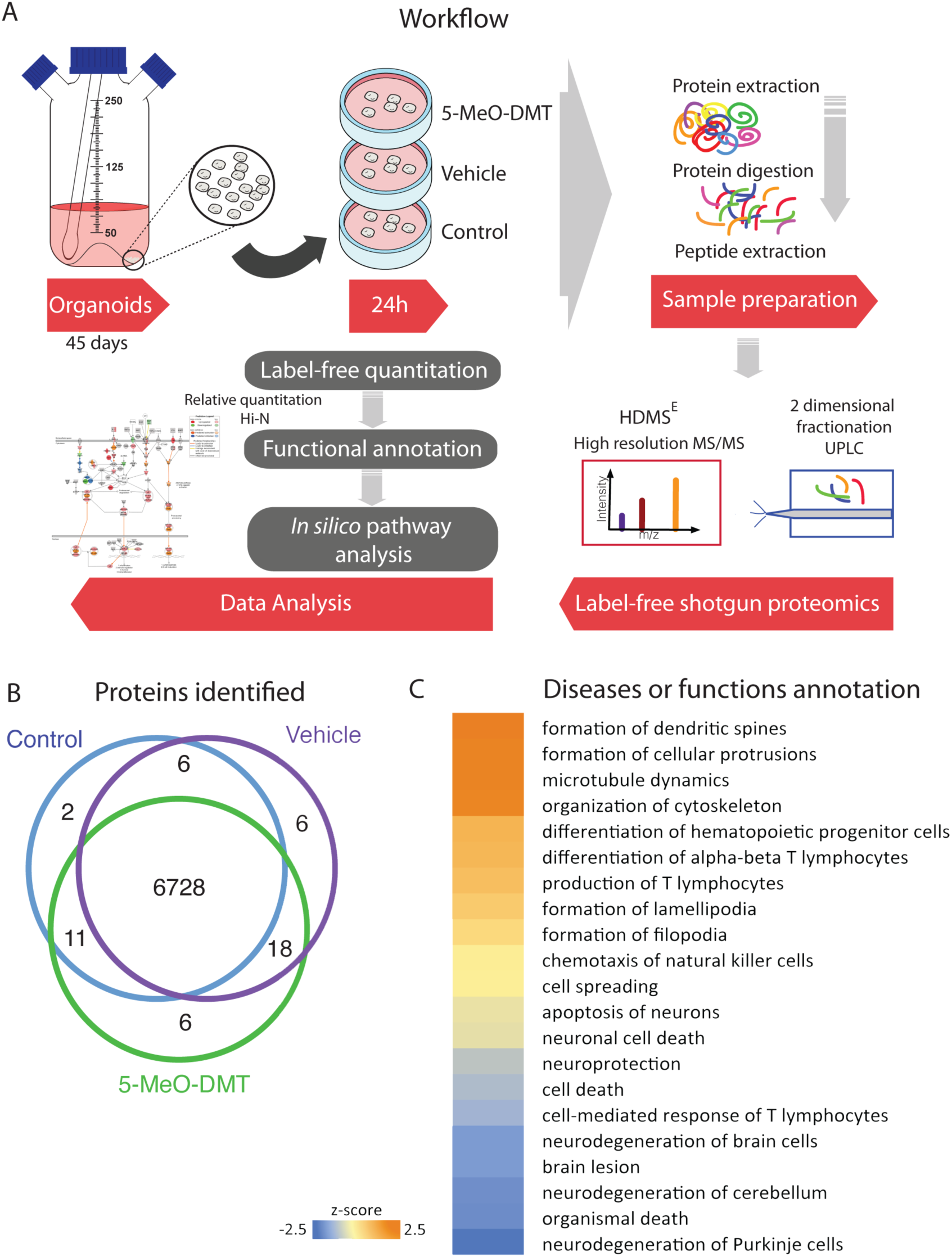
5-MeO-DMT treatment effects on human cerebral organoid proteomics. **(A)** Experimental design workflow. 45-day-old cerebral organoids were treated with 5-MeO-DMT, vehicle or untreated for 24h. Samples were analyzed using label-free state-of-the-art quantitative proteomics on a two-dimensional fractionation and high-resolution mass spectrometry. **(B)** Venn diagram comparing the number of proteins identified by shotgun mass spectrometry in control human cerebral organoids, and those treated with vehicle (EtOH), or 5-MeO-DMT. **(C)** Heat map showing significant functional enrichment between 5-MeO-DMT versus vehicle human cerebral organoids.

### 5-MeO-DMT leads to inhibition of NF-κB signaling pathway

Among the canonical pathways identified are nuclear factor of activated T-cells (NFAT) and nuclear factor kappa B (NF-κB) signaling via toll-like receptor (TLR), and Gq-coupled receptors, which are all inhibited by 5-MeO-DMT treatment (Fig. 4). Interestingly, the direct targets of 5-MeO-DMT, receptors 5-HT_2A_ and 5-HT_2C_, are Gq-coupled. Furthermore, NF-κB is very well known as the main transcriptional regulator of inflammatory, pro-inflammatory and anti-inflammatory cytokines and chemokines (Szabo & Rajnavolgyi, 2013).

**Figure 4.**
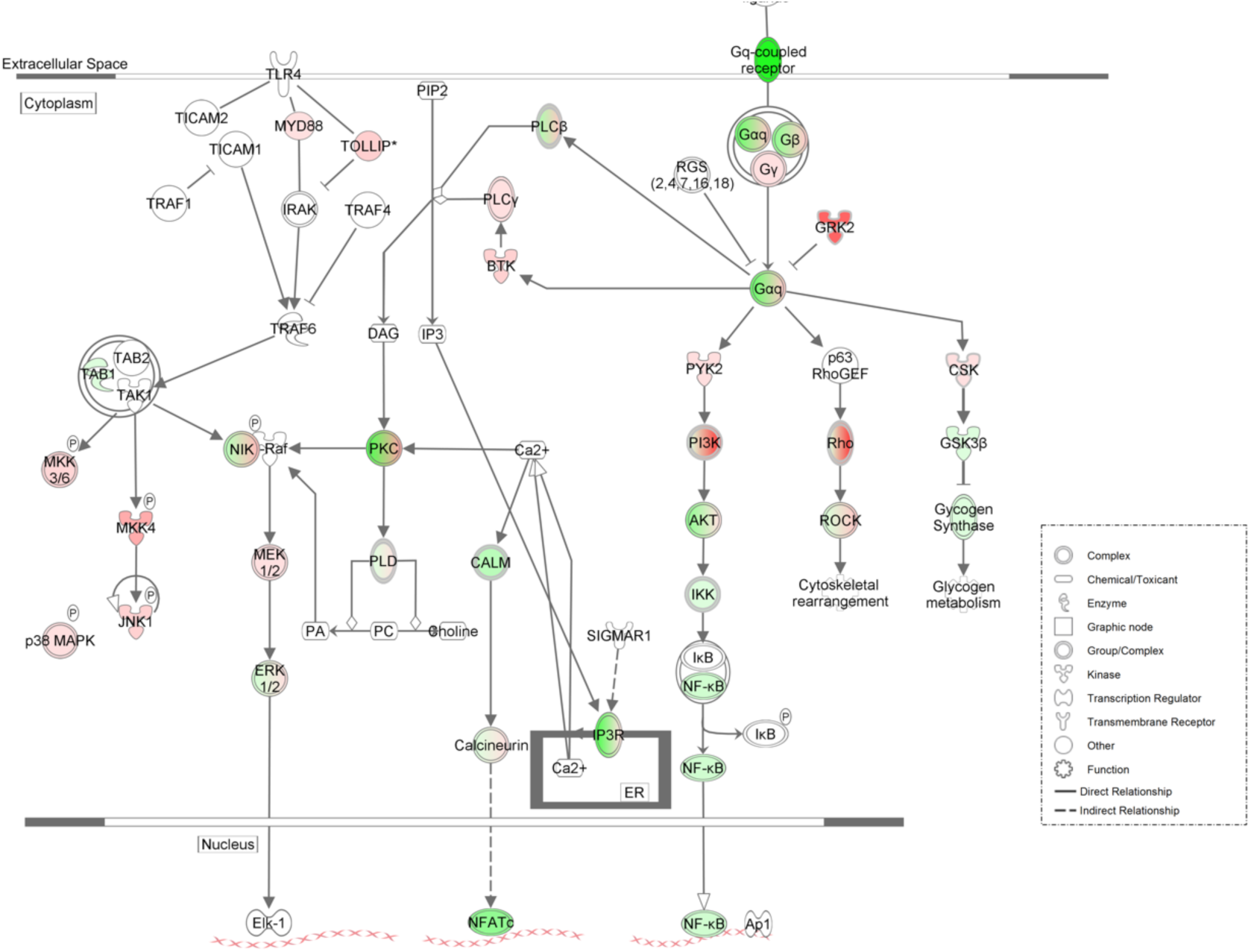
Schematic representation of the changes in protein expression of NFAT and NF-KB pathways by 5-MeO-DMT. Canonical pathways showing upregulated (red) and downregulated proteins (green) after 5-MeO-DMT treatment.

### Long-term potentiation components are modulated by 5-MeO-DMT

We have also identified regulation of specific proteins that participate in LTP, one of the main properties of most excitatory synapses throughout the CNS (Malenka and Bear, 2004). Proteins found upregulated are NMDAR (ionotropic glutamate receptor, NMDA), Ca^2+^/calmodulin-dependent protein kinase (CaMK2) and CREB (cyclic AMP-responsive element-binding protein). The group of downregulated proteins included mGluR5, G_αq_ protein, protein kinase C (PKC), PLC, calmodulin (CaM), AC1/8, inositol 1,4,5-trisphosphate receptor (IP3R), exchange factor directly activated by cAMP 1 (EPAC1) and PKA. These changes in key components, and further regulation of several other proteins and secondary messengers suggest a complex regulation of this pathway. AMPAR (AMPA-selective glutamate receptor 2), and the signaling cascade leading to c-Raf, mitogen-activated protein kinase kinase 1/2 (MEK1/2), extracellular regulated kinase 1/2 (ERK1/2) are upregulated, suggesting activation (Fig. 5). Based on the literature, activated ERK1/2 is transported to the nucleus and activates CREB, resulting in the expression of a large number of downstream genes (Alberini, 2009).

**Figure 5.**
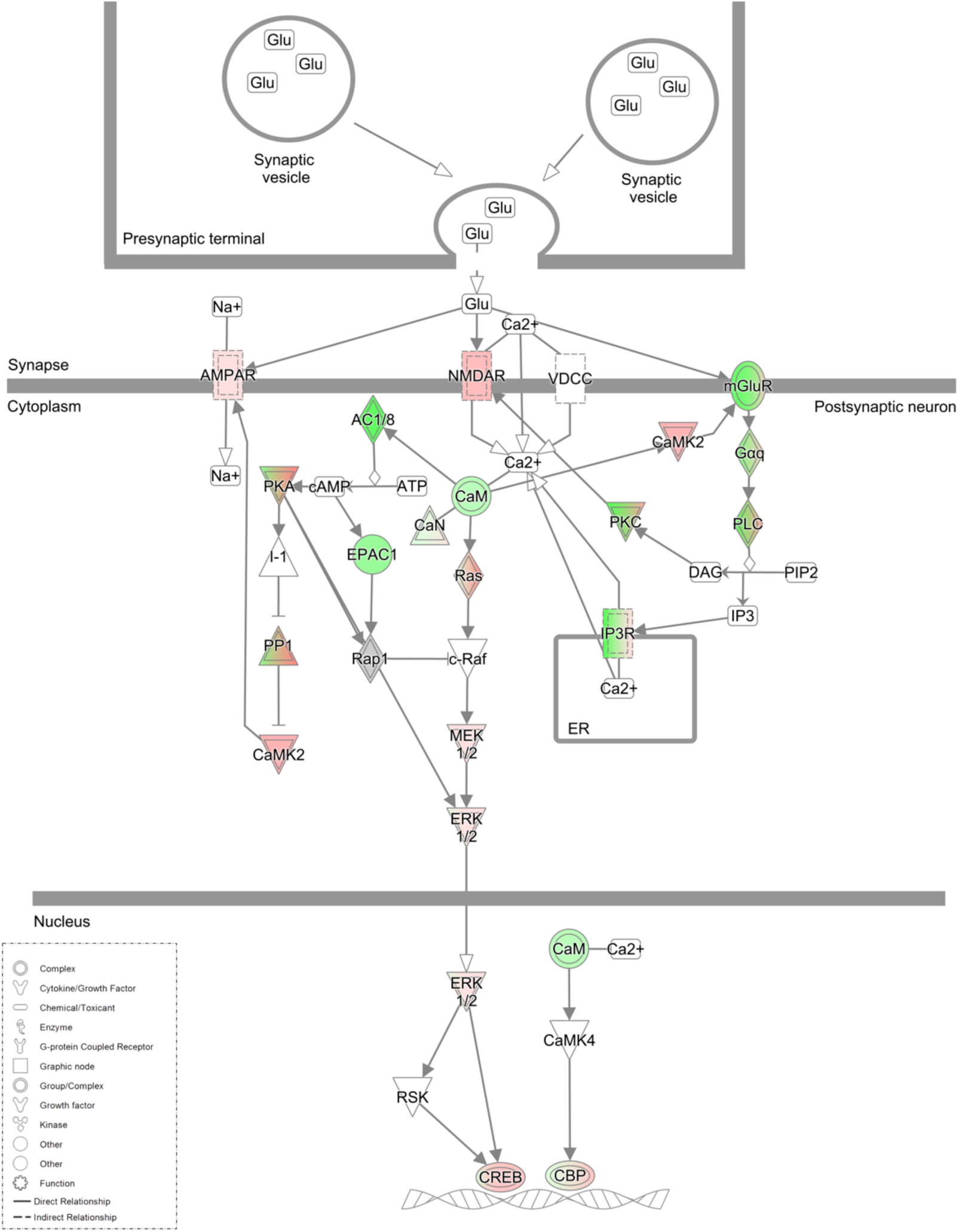
Schematic representation of proteins involved in long-term potentiation succeeding 5-MeO-DMT treatment. Z-scores were calculated from an upstream shortest path analysis and give the probability that the interaction between the proteins and the common regulator is not occurring by chance. In red, upregulated proteins; in green, downregulated proteins after 5-MeO-DMT treatment. Glu, glutamate.

### Cytoskeletal reorganization and dendritic spine morphogenesis proteins altered by 5-MeO-DMT

Ephrin B was another canonical pathway upregulated, including both forward and reverse signaling, as shown by the analysis of proteins differentially expressed (see Fig. 6). Upregulation of ephrin-B2 causes activation of ephrin type-B receptor (EPHB) and the cascade of CDC42 (cell division control protein 42 homolog), N-WASP (neural Wiskott-Aldrich syndrome protein) and ARP2/3 (actin-related protein 2/3) over intersectin, and interaction of ELMO1 activating RAC1 (Ras-related C3 botulinum toxin substrate 1), triggering dynamic reorganization of actin cytoskeleton and dendritic spine morphogenesis in forward signaling (Boyd *et al,* 2014). Meanwhile reverse signaling activates plexin, a protein that acts as a receptor for semaphorin over NCK adaptor protein 2 (GRB4) and focal adhesion kinase (FAK), causing axon repulsion through paxillin (PXN).

**Figure 6.**
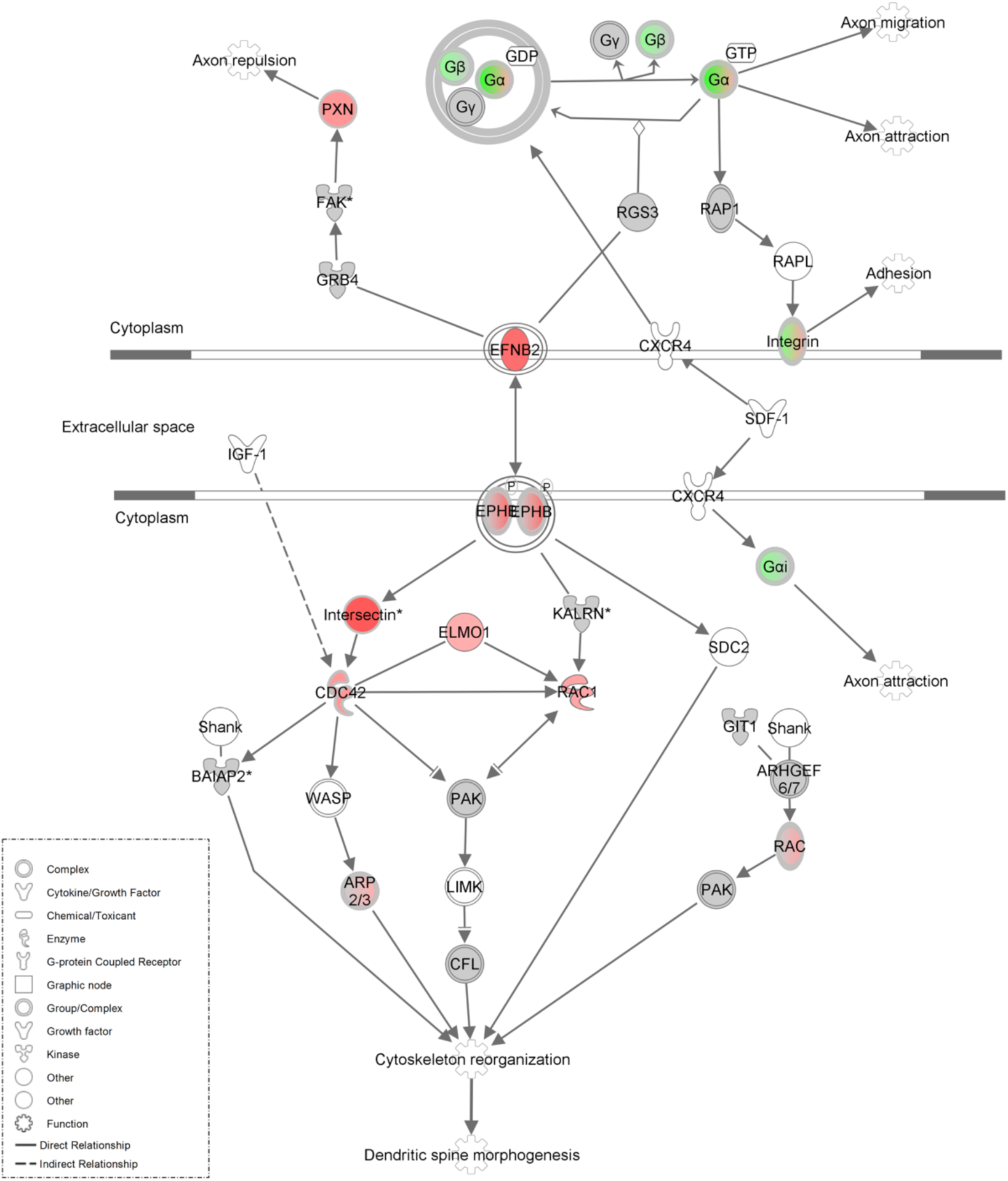
Pathway showing influence of 5-MeO-DMT on proteins engaged in cytoskeletal reorganization and dendritic spine morphogenesis. Canonical pathway showing upregulated (red) and downregulated proteins (green).

Additionally, we found significant regulation of plexins, integrins, SLIT-ROBO Rho GTPase-activating protein (srGAP), Netrin receptor DCC, metalloproteinase (Table 1) on 5-MeO-DMT-treated cerebral organoids, which corroborates actin regulation and orchestrate cytoskeletal reorganization.

**Table 1.**
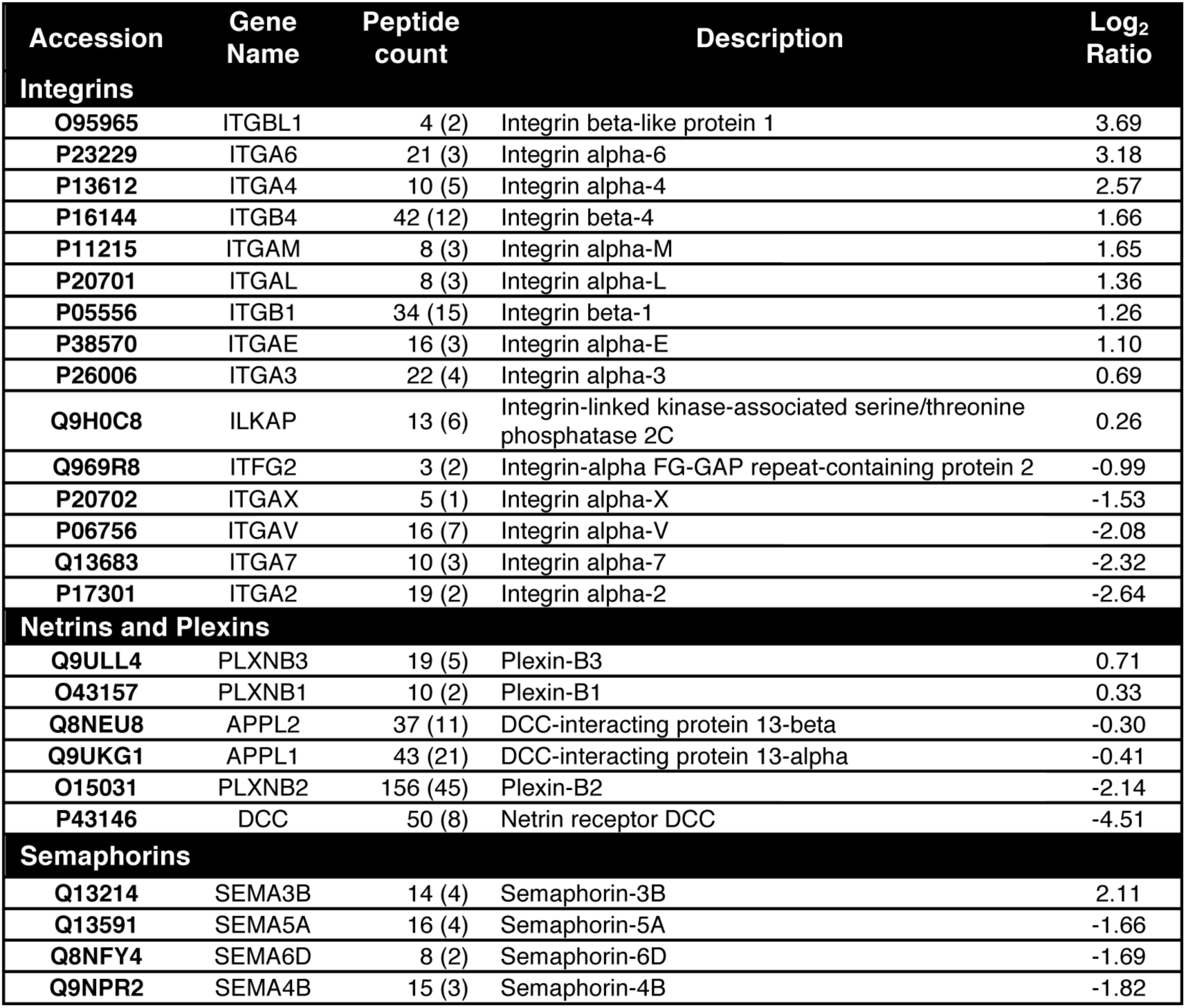
Table of proteins showing regulation of integrins, netrins, plexins and semaphorins by 5-MeO-DMT.

## Discussion

5-MeO-DMT is a structural analog of serotonin and melatonin and a functional analog of other psychedelic tryptamines such as N,N-DMT and 5-HO-DMT, a group of molecules we know very little about. The present results suggest that 5-MeO-DMT modulates the anti-inflammatory response, as well as the formation and maturation of novel dendritic spines, via proteins implicated with cellular protrusion formation, microtubule dynamics, cytoskeletal reorganization, and LTP. These changes were observed in the organoids and not in monolayer cultures of neuronal cells, which suggests a more mature and complex 3D circuitry are necessary for the actions of 5-MeO-DMT.

Here we demonstrate anti-inflammatory effects of 5-MeO-DMT using human cerebral organoids. NFAT and NF-κB signaling pathways were shown to be downregulated, via Toll-like receptors (TLR) and Gq-coupled protein receptors, most probably 5-HT_2A_ and 5-HT_2C_. Anti-inflammatory effects of 5-MeO-DMT were previously reported on human monocyte-derived dendritic cells, where inflammatory cytokine and chemokine release was shown to be blocked (Szabo *et al,* 2014). The immunomodulatory potential of other serotonergic psychedelics like lysergic acid diethylamide (LSD) (House *et al,* 1994; Voss and Winkelhake, 1974), 3,4-methylenedioxy-methamphetamine (MDMA) (Boyle and Connor, 2010; Connor *et al,* 2000), and 2,5-dimethoxy-4-iodoamphetamine (DOI) (Nau *et al,* 2013; Yu *et al,* 2008) were also previously reported. It is hypothesized that there is a cross-talk between TLR, serotonin receptors and sigma 1 receptors (Szabo, 2015).

Our work also revealed that only a 24h-treatment with 5-MeO-DMT, i.e., a single dose, modulates specific signaling molecules identified as key players in LTP, a classic mechanism of learning and memory (Malenka and Bear, 2004). Based on *in silico* predictions using proteomics data, modulation of these signaling molecules by 5-MeO-DMT would produce a complex regulation of LTP. One possibility is that LTP may be augmented in some cell types and inhibited in others, leading to a mixed protein signaling profile.

Additionally, we observed major downregulation of mGluR5 after treatment with 5-MeO-DMT, which has a role in the rewarding effects of several drugs of abuse. It was shown that mice lacking the mGluR5 gene do not self-administer cocaine and show no cocaine-induced hyperactivity (Chiamulera *et al,* 2001). They also have attenuated somatic signs of nicotine withdrawal (Stoker *et al,* 2012), and reduced ethanol consumption behavior (Bird *et al,* 2008), suggesting mGluR5 may be involved in addiction. The same effect has been demonstrated with the use of mGlu5 receptor antagonists for cocaine, nicotine, and ethanol in rats (Stoker *et al,* 2012). This effects of 5-MeO-DMT possibly can explain the therapeutic effect of *Ayahuasca* on substance dependence (Barbosa *et al,* 2012; Crews *et al,* 2011; Doering-Silveira *et al,* 2005; Fábregas *et al,* 2010; McKenna, 2004; Thomas *et al,* 2013). Moreover, *Ayahuasca* seems to inhibit addictive behaviors in an animal model of alcohol dependence (Oliveira-Lima *et al,* 2015). In humans, Ayahuasca administration to healthy subjects reduces rapid-eye-movement sleep (REM) and increases slow-wave sleep (SWS) (Barbanoj *et al,* 2008), two major sleep stages respectively associated with increased or decreased LTP (Blanco *et al,* 2015; Dumoulin Bridi *et al,* 2015; Ribeiro, 2012; Tononi and Cirelli, 2014), while producing dream-like effects. Thus, the plausible complex regulation of LTP-related proteins detected here also may indicate that 5-MeO-DMT produces a mix of SWS and REM-like effects.

Changes in LTP/LTD balance are directly connected with increase/decrease of dendritic spine number and size, respectively (Bourne and Harris, 2008). Spine morphogenesis relies on alterations in the actin cytoskeleton, but the molecular mechanisms that regulate this process are still not clear. 5-MeO-DMT caused significant upregulation of EFNB2, EPHB, and various secondary messengers involved in dendritic spine formation. Dendritic spine formation can be caused by direct stimulation of serotonergic receptors. Indeed the selective 5-HT2A receptor agonist, DOI, have been shown to modulate spine morphology of mature cortical pyramidal neurons (Jones *et al,* 2009). In this study, a transient increase in spine size induced by DOI was kalirin-dependent and enhanced phosphorylation of PAK, whereas here we show upregulation of EFNB2, EPHB, intersectin, ELMO1, CDC42 and RAC1. Binding of ephrin-Bs to EphB receptors initiate bidirectional signaling, which, by altering actin cytoskeleton, leads to changes in dendritic spine shape, size, and number (Klein, 2009). It was shown that EPHB2 interacts with intersectin and activates its GEF activity in cooperation with N-WASP, which in succession activates the Rho-family GTPase Cdc42 and spine morphogenesis (Irie and Yamaguchi, 2002). N-WASP is a critical regulator of Arp2/3-mediated actin polymerization (Takenawa and Miki, 2001). Henkemeyer and colleagues demonstrated that triple EphB1,2,3-deficient hippocampal neurons have abnormal formation of actin clusters along dendrites, impairing normal dendritic spine formation *in vivo* (Henkemeyer *et al,* 2003). Meanwhile, *in vitro,* knockdown of EphB2 alone is sufficient to reduce synapse density (Kayser *et al,* 2006). Postnatal re-expression of EphB2 in slice cultures from animals lacking EphB1–3 is sufficient to rescue dendritic spine defects (Kayser *et al,* 2006). Although EphB signaling has a clear role in dendritic spine morphogenesis through kinase domain activity, it can also regulate activity-dependent synaptic plasticity interacting with both NMDA (Takasu *et al,* 2002) and AMPA receptors (Kayser *et al,* 2006). Literature shows that σ-1R also could contribute to the brain plasticity effects of 5-MeO-DMT. σ-1 R is an endogenous regulator of dendritic spine morphology (Ruscher *et al,* 2011; Tsai *et al,* 2009) and neurite outgrowth (Ruscher *et al,* 2011).

Apart from acting as a direct molecular mediator of plasticity, 5-MeO-DMT had effects on cell surface and extracellular proteins involved in regulating synaptic architecture, like plexins (Laht *et al,* 2014), DCC (Horn *et al,* 2013), metalloproteinase (Bozdagi *et al,* 2007) and integrins (Dityatev and Schachner, 2006; Shi and Ethell, 2006). An upregulation of integrins, as we observed here in 5-MeO-DMT-treated organoids, was also found in major depression patients who presented good response to antidepressants, suggesting the importance of this class of proteins in brain plasticity (Martins-de-Souza *et al,* 2014). One more protein significantly downregulated is srGAP, an intracellular signaling molecule with a role in processes underlying synaptic plasticity, higher cognitive function, learning, and memory (Endris *et al,* 2002).

Finally, we also found important functions such as neurodegeneration, cell death, and brain lesion inhibited by 5-MeO-DMT. These neurorestorative and cellular protective effects are expected after activation of σ1R (Frecska *et al,* 2013; Szabo *et al,* 2016). σ1R agonists exert neuroprotective effects by regulating intracellular calcium levels (Mueller *et al,* 2013), preventing expression of pro-apoptotic genes (Tchedre and Yorio, 2008) and protecting mRNA of anti-apoptotic genes, such as Bcl-2.

Fast antidepressants also have strong effect on synaptic plasticity, reversing functional and structural synaptic deficits caused by stress. A typical example of this group is ketamine, an hallucinogenic, non-competitive NMDA glutamate receptor channel antagonist, which causes an improvement in mood ratings within hours, as opposed to weeks as in typical antidepressants (Duman *et al,* 2016). Ketamine increases mammalian target of rapamycin complex 1 (mTORC1) signaling, via activation of Protein kinase B (PKB or Akt) and ERK. mTOR signaling then boosts synaptic protein synthesis, spine stability and function in the prefrontal cortex (Duman *et al,* 2016; Duman and Aghajanian, 2012; Gerhard *et al,* 2016).

Therefore, the pattern of proteins altered after 5-MeO-DMT treatment points to robust actions on synaptic plasticity and improvement of cell survival. Taken together our data offer a possible mechanistic insight into the neural changes produced by the chronic ingestion of substances containing dimethyltryptamines.

## Acknowledgements

This work is part of the PhD thesis of VD. We thank Yury M. Lages, Dr. Sylvie Devalle, Ismael Gomes, Marcelo Costa and Gabriela Vitoria for technical assistance. We also would like to thank Dr. Richardson Leão, Dr. Marília Zaluar Passos Guimarães and Dr. Eduardo Schenberg for support during this project.

## Author Contributions

VD, JMN, SR designed the experiments; VD, JMN, RMM, RS performed the experiments; DMS supervised analysis and interpretation of the proteomics data; VD, JMN, and RMM analyzed the data; VD, JMN, SR wrote the paper; VD, JMN, RMM, RS prepared figures; VD, JMN, RMM, RS, DA, SR, DMS and SR reviewed drafts of the paper.

